# Five years later, with double the demographic data, naked mole-rat mortality rates continue to defy Gompertzian laws by not increasing with age

**DOI:** 10.1101/2023.03.27.534424

**Authors:** J. Graham Ruby, Megan Smith, Rochelle Buffenstein

## Abstract

The naked mole-rat (*Heterocephalus glaber*) is a mouse-sized rodent species, notable for its eusociality and long lifespan. Previously, we reported that demographic aging, i.e., the exponential increase of mortality hazard that accompanies advancing age in mammals and other organisms, does not occur in naked mole-rats (Ruby et al, 2018). The demographic data supporting that conclusion had taken over three decades to accumulate, starting with the original rearing of *H.glaber* in captivity. In the five years following that study, we ∼doubled our quantity of demographic data. Here, we re-evaluated our prior conclusions in light of these new data and found them to be supported and indeed strengthened. We additionally provided insight into the social dynamics of captive *H.glaber* with data and analyses of body weight and colony size versus mortality. Finally, we provide a phylogenetically-proximal comparator in the form of lifespan data from our Damaraland mole-rat (*Fukomys damarensis*) colony and demographic meta-analysis of those data along with published data from Ansell’s mole-rat (*Fukomys anselli*). We found *Fukomys* mortality hazard to increase gradually with age, an observation with implications on the evolution of exceptional lifespan among mole-rats and the ecological factors that may have accompanied that evolution.

## Introduction

The species *Heterocephalus glaber*, commonly known as the naked mole-rat, is a eusocial mammal endemic to the arid and semi-arid regions of northeast Africa. In the wild, naked mole-rats live an almost completely subterranean lifestyle, in colonies of up to 295 animals (average size is 60 animals/colony; Brett, 1991) that cohabitate a network of tunnels that the mole-rats dig themselves with their large, ever-growing incisors (Buffenstein et al, 2021). Each colony is usually led by a single, socially-dominant queen (breeding female), living with a small number of breeding males and a large number of non-breeding progeny. New breeding females arise either through conflict following the death of a queen or isolation of part of the colony, e.g. through tunnel collapse (Brett, 1991; Jarvis, 1981; Buffenstein et al., 2022).

Naked mole-rats are notable for their extreme lifespans (Buffenstein 2005; Lewis and Buffenstein 2016), living longer than any other documented rodent (Buffenstein and Jarvis, 2002), with the longest previously-reported lifespan of 37 years (Lee et al.,2020) and many animals living beyond 30 years (Lewis et al., 2018; Can et al., 2022). These values are notable in the context of this species’ small body size due to the strong correlation across species between that value and mammalian lifespan: maximum lifespan potential (MLSP) increases by 16% for each doubling of average species body mass (Hulbert et al., 2007). Notably, naked mole-rats live approximately five times longer than their size-corrected, allometric expectation using the equations of (de Magalhães et al., 2007). For most mammalian species, lifespan is limited by an exponential increase in the per-day risk of death (i.e. mortality hazard) with age (Gavrilova & Gavrilov, 2015; Cohen, 2018) in accord with a statistical distribution first defined by Gompertz based on human mortality (Gompertz, 1825). The increase in mortality hazard with age is referred to as “demographic aging”. We have previously shown *H.glaber* to achieve its exceptional longevity through defiance of this trend, exhibiting ∼constant mortality hazard across the full spectrum of observed lifespans, with no hazard increase evident even many-fold beyond their expected MLSP (Ruby et al, 2018 & 2019).

Naked mole-rats are members of the the African mole-rat superfamily, Bathyergidae, that includes more than 30 species of subterranean rodents and appeared to split from the other bathyergids ∼26 million years ago (MYA) (Fang et al., 2014; Visser et al., 2019; Patterson and Upham, 2014; Faulkes and Bennett 2021). Bathyergid species span the spectrum of social behavior, from solitary (e.g., *Georychus, Heliophobus* and *Bathyergus*) to the only other known eusocial mammals, members of the genus *Fukomys* (Jarvis and Bennett, 1993). Along with eusociality, *H.glaber* also share above-expected longevity with members of the *Fukomys* genus, namely *Fukomys damarensis* and *Fukomys anselli,* living ∼twice as long as their size-corrected, allometric expectations (see Methods for details). The co-occurrence of these unusual properties in divergent mole-rat clades motivates a question about the similarity of age-related hazard patterns.

For naked mole-rats, the lack of demographic aging is accompanied by seemingly-indefinite maintenance of many physiological characteristics that typically change with age. Naked mole-rats are resistant to age-related diseases such as cancer, neurodegeneration, and cardiovascular disease (Edrey et al., 2011; Hadi et al., 2021; Can et al., 2022) and show signs of tissue regeneration and remodeling preventing the deterioration of age-associated physiological function (Buffenstein et al., 2020, Can et al., 2022; O’Connor et al., 2002; Buffenstein and Craft, 2021). However, controversy regarding the perpetual neoteny of *H.glaber* (Orr et al., 2016; Grimes et al., 2017; Skulachev et al., 2017; Buffenstein et al., 2020) remains, supported by evidence for chronological aging of the epigenetic (DNA methylation) clock, albeit showing that breeders maintain a more youthful methyl clock (Kerepesi et al., 2022). This is further stoked by a commentary on our original demographic analysis (Ruby et al, 2018) by (Dammann et al, 2019), asserting that loss of records from our early years of our vivarium rearing of *H.glaber* could have confounded our analysis of aging demographics. The concern of that critique has been addressed using left-censorship (Ruby et al, 2019). However, the critique also highlights the unique nature of our collection as a resource for powered demographic analysis of naked mole-rats, extending out to advanced ages. The lack of other collections of a similar scale, worldwide, presents a practical challenge to the reproducibility of our results. To that end, we were motivated to re-evaluate our original conclusions, using demographic data for *H.glaber* collected over a five-year period since the data freeze from (Ruby et al, 2018).

Here, we re-visited the demographic analysis of our naked mole-rat collection, with husbandry data now extended by five years across an expanded set of animals. We found our original conclusions of naked mole-rat mortality hazard being age-independent to be reproduced, using either the total sum of all historical data or only those data collected after our previous study – the latter qualifying as a replication study. We further conducted an analysis of naked mole-rat mortality hazard versus colony size which suggested that social competition was greater for non-breeders in smaller colonies. Finally, we provide demographic data for Damaraland mole-rats: combining those data with other publicly available data for *Fukomys* species, we analyzed age-specific mortality hazard and showed it to increase, in contrast to the naked mole-rats.

## Results

### The expanded data set revealed that the hazard of naked mole-rat mortality did not increase as a function of age

Our expanded data set drew from 7,536 historically catalogued naked mole-rats from our own collection; of these, 6,957 had sufficiently high (single-day) resolution records for date-of-birth and date-of-death/censorship to be included (see Methods for details). Our analyses of survival and mortality hazard began at T_sex_ for naked mole-rats: 6 months (183 days; Edrey et al., 2011); 6893 animals had aged up to T_sex_ and therefore were included in our downstream analyses. Of those, 755 had recorded deaths and 6138 were right-censored (see Methods for censorship criteria). Kaplan-Meier survival, along with death and censorship distributions, are plotted in Figure 1A. The last recorded death was at 11,281 days-of-life (30.9 years), at which point 10 animals remained alive; the last of those was censored at 12,873 days-of-life (35.2 years). At that event, Kaplan-Meier survival remained at 55.5%.

**Fig 1:**
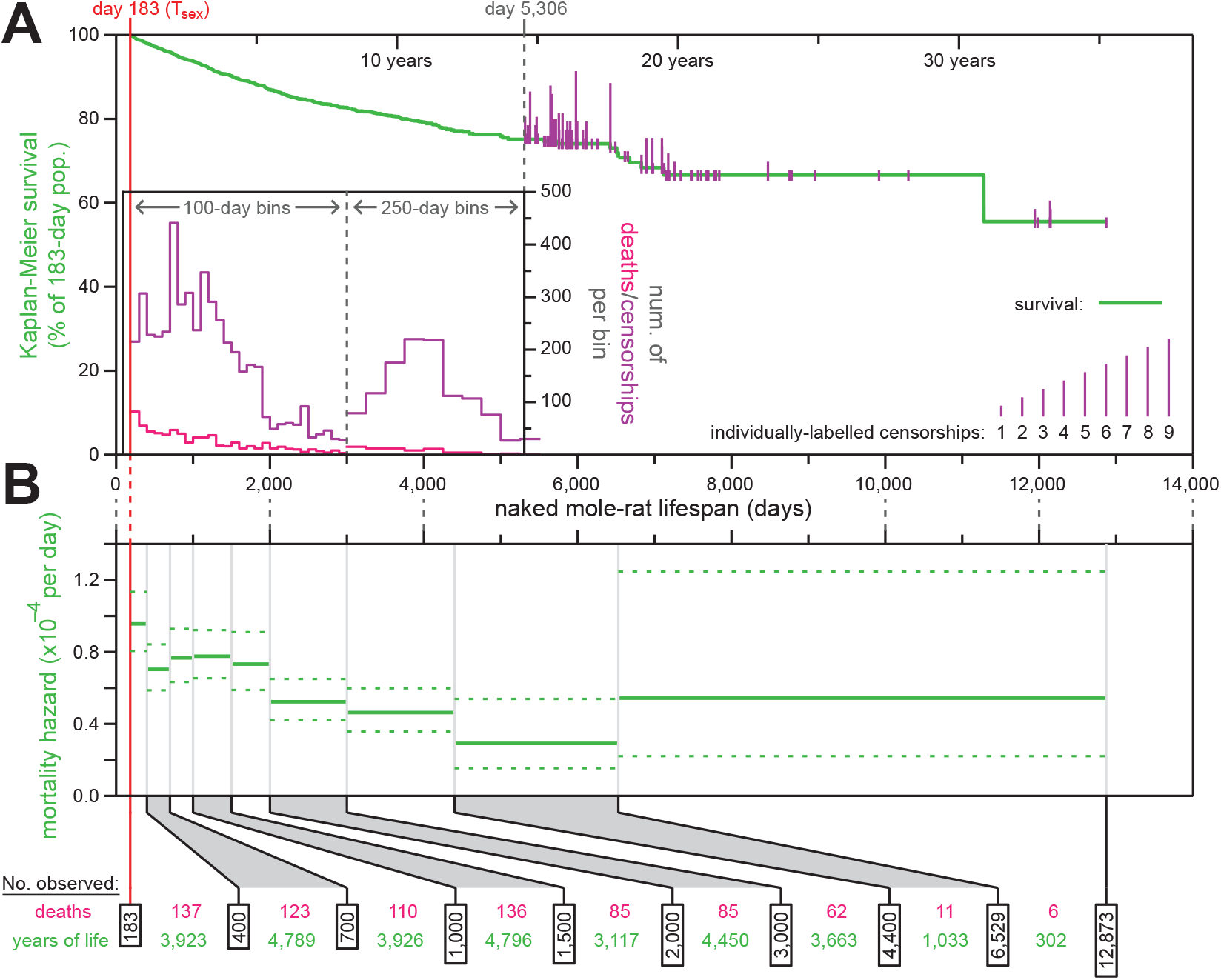
The mortality hazard of naked mole-rats failed to increase with age. (A) Kaplan-Meier survival curve for naked mole-rats (green) after reaching reproductive maturity (T_sex_; 6 months from birth; 183 days; red). Plotted as in Fig. 1A of (Ruby et al., 2018), with a histogram of death and censorship (pink and purple, respectively) in the inset, up until 5,306 days of age, at which point 174 animals remained in the population; and with censorship events after that indicated by vertical ticks (purple), the size of which is proportional to the number of animals censored at each day-of-life. (B) Mortality-hazard estimates (solid green) with 95% confidence intervals (dotted green) across the observed naked mole-rat lifespan. Estimates were calculated across time intervals of increasing size (demarked by grey lines) to compensate for decreasing accuracy-per-unit-time as the population size decreased. The borders of the age bins (black), along with the number of deaths (pink) and total years of animal life observed (green) per bin, are indicated at the bottom.

Using the survival data, we calculated age-specific mortality hazard across the same age-range windows as in (Ruby et al, 2018) (Figure 1B). As in the prior analysis, mortality hazard estimates remained consistently low across the ∼35 years of age encompassed by the observations, never exceeding a per-day probability of 1/10,000.

### Analysis of only data collected after our prior survey independently confirmed constant-hazard demography

The large size of our naked mole-rat collection across the five years of additional observation time since our previous report (Ruby et al, 2018) provided substantial additional power in the form of number of animals observed at most ages (Fig. 2A). For many age groups, the amount of lifespan observation time since our previous snapshot matched or exceeded that prior to it. This allowed us to use left-censorship at a date after our original dates of record aggregation to perform an independent analysis of naked mole-rat lifespan demographics and critically re-evaluate our original conclusions.

**Fig 2:**
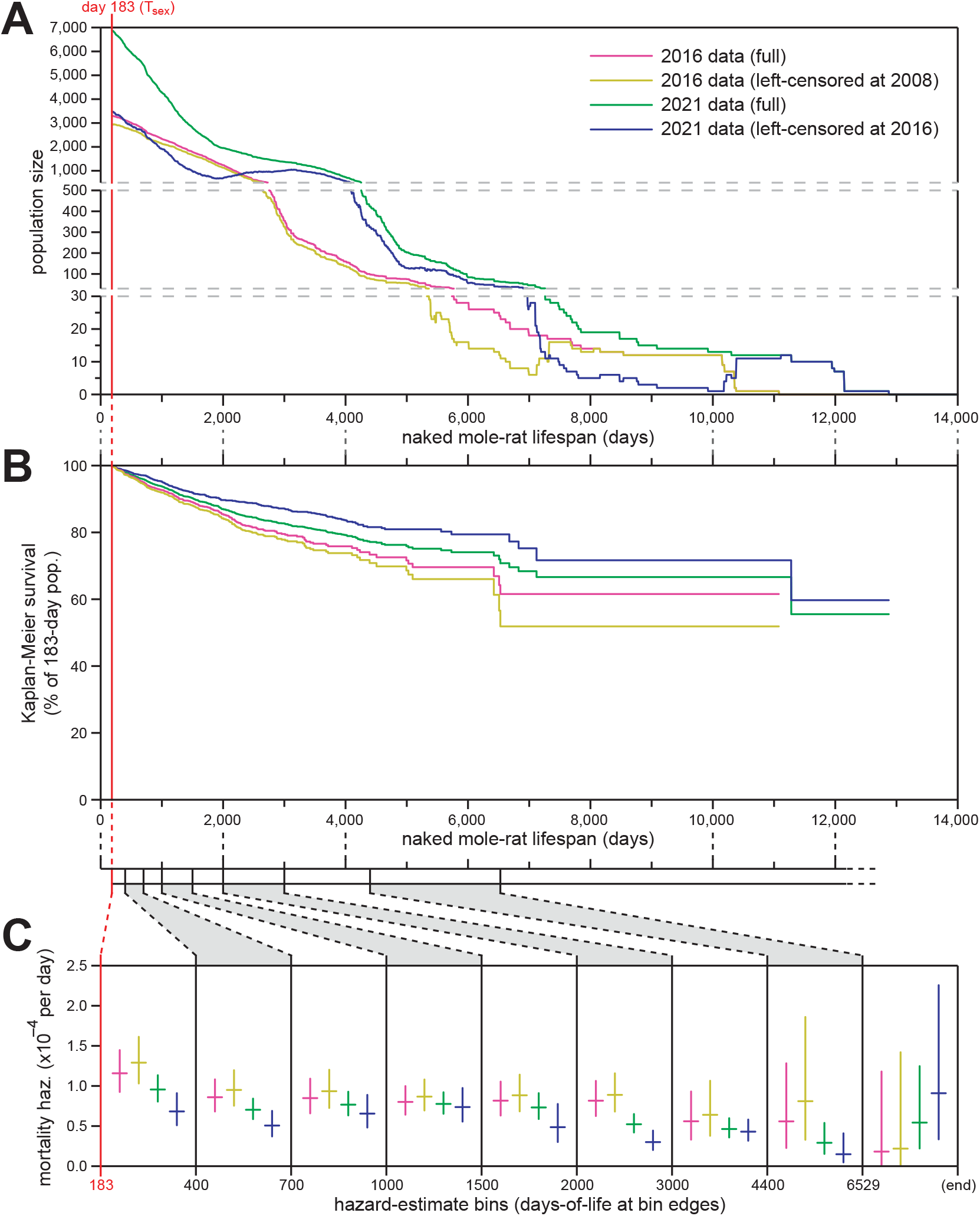
Exclusion of neither pre-2008 nor pre-2016 lifespan data through left-censorship modified the observed lifespan demographics of *H.glaber*. (A) The number of naked mole-rats observed (y-axis) at each age (x-axis) for four datasets: i) the original data set presented in (Ruby et al., 2018) (pink); ii) that data set left-censored on January 1, 2008, as presented in (Ruby et al., 2019) (yellow); iii) the full data set compiled in 2021 for this manuscript (green); iv) that data set left-censored on May 17, 2016 (navy), i.e. data collected after the compilation of data for (Ruby et al., 2018). (B) Kaplan–Meier survival curves starting at T_sex_ (183 days; red) for the four demographic data sets, as colored and described in panel (A). (C) Mortality hazard estimates (y-axis) across each of the lifespan bins from Figure 1B (indicated on the x-axis), for each of the four demographic data sets, as colored and described in panel (A). Horizontal bars indicate hazard estimates, and vertical bars indicate 95% confidence intervals.

For our previous report, lifespan records were aggregated in three batches: on April 14, 2016; April 20, 2016; and May 17, 2016 (Ruby et al, 2018). Animal counts as a function of age, Kaplan-Meier survival, and age bin-specific hazard are shown for that data set in magenta in Fig. 2A, B, and C, respectively. The possibility that incomplete record survival during the earliest years of our keeping naked mole-rats in captivity could have skewed mortality hazard trends was raised by Dammann et al (2019), and we previously addressed that concern by excluding data from those early years of naked mole-rat captivity, via left-censorship on January 1, 2008 (Ruby et al, 2019). No additional data were provided at that time. The scale, Kaplan-Meier survival, and age-specific hazard from that left-censored version of our original data set are shown in orange in Fig. 2A, B, and C, respectively.

The scale, Kaplan-Meier survival, and age-specific hazard from our current, full data set – also depicted in Figure 1 – are shown in green in Figures 2A, 2B, and 2C, respectively. These data were aggregated on March 15, 2021 and encapsulated all historical data. In order to perform an analysis of naked mole-rat lifespan demographics that would be independent of our original analysis, we left-censored these data on May 17, 2016. The scale, Kaplan-Meier survival, and age-specific hazard from this left-censored data set are shown in blue in Fig. 2A, B, and C, respectively. For each age bin, the new, left-censored hazards were either lower or statistically indistinguishable from the original data set, either full or left-censored (Fig. 2C). We therefore maintain our original conclusion: that naked mole-rat mortality hazard does not increase with age, with that assertion being statistically robust to the beginning of our final estimate bin (6,529 days, or ∼18 years).

### Non-increasing mortality hazard was exhibited by breeders and non-breeders of both sexes

The analyses described above were performed on our full population of animals. Sub-populations defined by sex and breeding-status were analyzed individually for Kaplan-Meier survival (Figure 3A). As previously observed (Ruby et al, 2018), there was a notable survival difference between breeding versus non-breeding status, with breeders enjoying longer survival than non-breeders. As with the aggregate population, mortality hazard remained consistent with age, for both non-breeders (Figure 3B) and breeders (Figure 3C), with the mortality hazard of breeders only ∼20% that of non-breeders. As previously observed, male non-breeders exhibited enhanced survival (Figure 3A) and reduced mortality hazard (Figure 3B) at ages beyond 2000 days. This concurred with the expectations of our model that multiple males can enjoy breeder status in a colony despite not being labeled as such (only the founding breeders are so labeled; Ruby et al, 2018).

**Fig. 3.**
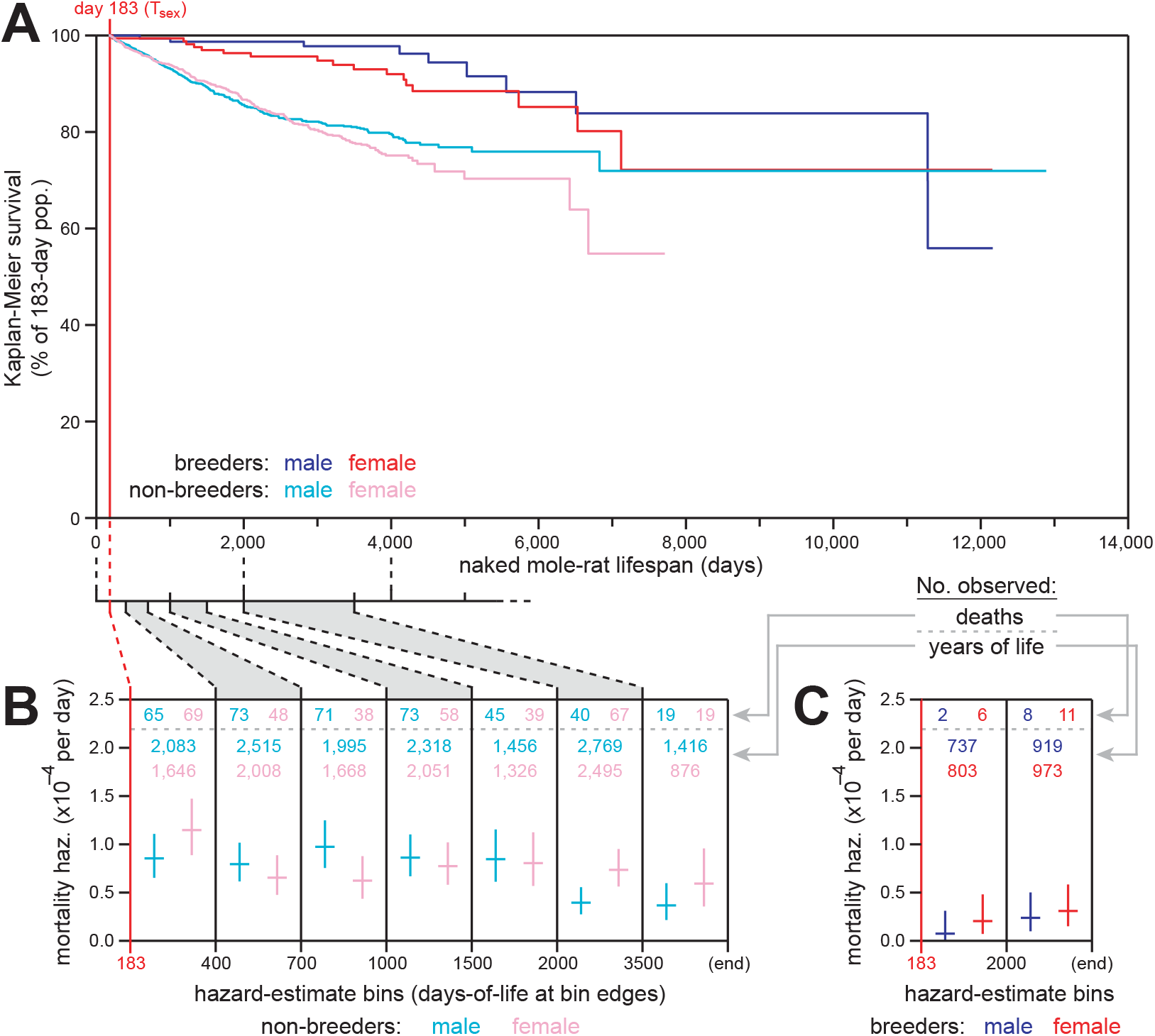
Breeding and non-breeding male and female naked mole-rats exhibited non-increasing mortal hazard as a function of age. (A) Kaplan-Meier survival starting at T_sex_ (183 days; red) for each of four reproductive categories of naked mole-rats: male breeders (navy), female breeders, (red), male non-breeders (cyan), and female non-breeders (pink). (B) Mortality hazard estimates (y-axis) for the indicated lifespan bins (x-axis), calculated for non-breeding males (cyan) and females (pink). The numbers of observed death events per bin are indicated at the top of each bin. (C) Mortality hazard estimates (y-axis) for the indicated lifespan bins (x-axis), calculated for breeding males (navy) and females (red). The numbers of observed death events per bin are indicated at the top of each bin.

### Non-breeder mortality hazard was influenced by colony size

Some of the mortality hazard difference between breeders and non-breeders may derive from social struggle between these hierarchies within the colony. To explore the potential effects of intra-colony social dynamics on mortality hazard, we performed analyses of the relationships between body weight or colony size on the mortality of non-breeders.

For body weights, we analyzed a data set that had been aggregated to accompany our originally published demographic data (Ruby et al, 2018; Supplementary Table 2). These data included 19,118 individual body weight estimates from 1,940 animals, captured longitudinally for animals of a variety of ages, with data becoming sparse after ∼3500 days of age (Figure 4A). These data are provided in Supplemental Data File 2. A great deal of variation existed across both the breeder and non-breeder populations, in both males and females, though the age-specific averages were similar and stable after ∼500 days of age (Figure 4B).

**Fig. 4.**
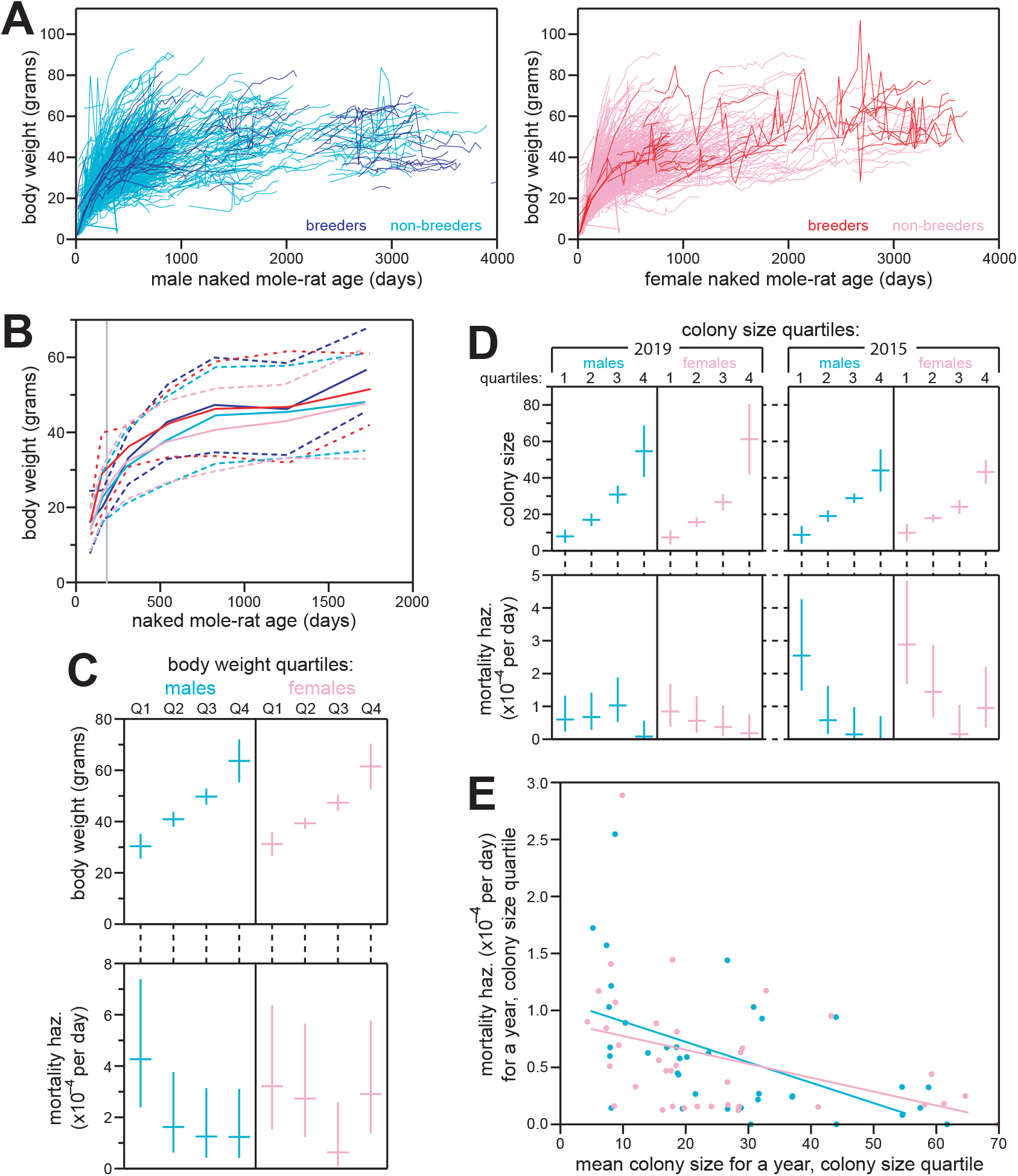
Non-breeding naked mole-rat mortality hazard decreased with increasing body weight and colony size. (A) Individual body weight values (y-axis) plotted longitudinally versus age (x-axis) for breeding males (navy), non-breeding males (cyan), breeding females (red), and non-breeding females (pink). (B) Mean body-weights for each of the categories from panel (A), for all weight measurements taken across each of the following age bins: 50-120 days, 120-200 days, 200-400 days, 400-700 days, 700-1000 days, 1000-1500 days, and 1500-2000 days; inclusive of the 1^st^ day and non-inclusive of the last day for each bin. Solid lines indicate mean body weights; dotted lines indicate standard deviations. (C) Top panel: body weight quartiles for male (left) and female (right) non-breeding animals, organized based on the last recorded weight measurement for each animal and only included if that measurement was taken at an older age than 500 days. Horizontal bars indicate the mean body weight for each quartile; vertical bars indicate standard deviations. Bottom panel: mortality hazard for each body weight quartile in the top panel, calculated for the year following weight measurement. See Methods for details. Horizontal bars indicate mortality hazard estimates; vertical bars indicate 95% confidence intervals. (D) Examples of colony-size versus mortality-hazard plots. For each indicated year (2019 on the left; 2015 on the right), and for each indicated non-breeding sex (males on the left; females on the right): animals were organized into quartiles based on the number same-sex, non-breeding animals in their colony. Top panels: the mean colony sizes for animals in each quartile; vertical bars indicate standard deviations. Bottom panels: mortality hazard estimates for each colony-size quartile from the top panel; vertical bars indicate 95% confidence intervals. Similar analyses were performed for each year, 2012-2020; all plots are provided in Supplemental Figure S1. See Methods for details. (E) For all colony-size quartiles, from all years analyzed in Supplemental Figure S1: a meta-analysis of quartile mean colony sizes (x-axis) versus one-year mortality hazards (y-axis). Non-breeding males (cyan) and females (pink) were meta-analyzed separately. Regression lines for each sex are shown (p-values: 9.8 * 10^-4^ for males; 0.018 for females).

For the analysis of mortality hazard versus body weight, only non-breeding animals were included. The last available body weight measurement was used, requiring that measurement was taken after 500 days of age, to avoid confounding with the social status of animals that were not yet fully grown. There were 6,785 qualifying measurements, derived from 942 animals. For each animal, mortality hazard was analyzed across one year following that final measurement. Non-breeding animals of each sex were grouped into quartiles based on body weight, and the aggregate mortality hazard of each quartile is plotted, along with the aggregate body weight values, in Figure 4C. There was a general trend of lighter/smaller animals having greater mortality hazard, especially in males, though the data set was insufficiently powered to statistically support this conclusion.

For colony sizes, we used our records’ annotations of “colony ID” to determine the number of non-breeding animals of each sex in each colony, at each analytical time point (see Methods). As for body weight, animals were separated into quartiles based on the number of non-breeders of the same sex in the colony, and each quartile for each sex was cumulatively evaluated for mortality hazard across one year following the analytical time point. This method allowed independent analysis of non-overlapping time periods: we performed five such analyses using our new data set and four using our prior data set, from Ruby et al, 2018 (examples in Figure 4D; all results shown in Supplemental Figure S2). Analyses of individual time blocks were statistically inconclusive, but there was a general trend of greater mortality hazard for non-breeders in smaller colonies. Meta-analysis of the results from across all nine time-block analyses supported this trend to a statistically significant degree, for both males (p-value: 9.9 * 10^-4^) and, to a lesser degree, females (p-value: 1.8 * 10^-2^).

### The mortality hazard of mole-rats of the Fukomys genus increased substantially with age

We have previously placed the mortality demographics of naked mole-rats into the context of demographic data from other mammals, which highlights the unique absence of an exponential increase in hazard with age in naked mole-rats (Ruby et al, 2018). To provide a more phylogenetically-proximal comparator, we performed a demographic analysis of mole-rats from the *Fukomys* genus. For those species, survival analyses all began at an estimated T_sex_ of 270 days of age, based upon our unpublished data and personal communication from Nigel C. Bennett (MRI, University of Pretoria) and Christopher G. Faulkes (Queen Mary University of London).

In their response to our previous publication, Dammann et al (2019) provide lifespan data for 339 *Fukomys anselli* individuals, of which 332 aged past T_sex_ and therefore were included in our downstream analyses (Supplemental Figure S2A: yellow). In their analyses of lifespan, Dammann et al (2019) censor accidental deaths and deaths from fighting: in that case, 170 animals’ lifespans are censored and 162 died. Here and previously (Ruby et al 2018,2019), we treat such instances as non-censored deaths: in that case, 145 animals’ lifespans are censored and 187 had died. Comparing Kaplan-Meier survival analyses, the death versus censorship of these 25 individuals had a small effect (Supplemental Figure S2B: yellow versus orange), only reducing the median lifespan from 3,141 to 2,830 days; nor were any statistical differences observed for age-specific mortality hazard estimates effect (Supplemental Figure S2C: yellow versus orange). For downstream analyses, we considered those 25 individuals to be dead rather than censored.

We provide additional demographic data for another mole-rat species within the *Fukomys* genus (*Fukomys damarensis*). These data were compiled on March 15, 2021 and included lifespans for 702 animals, of which 634 aged past T_sex_ and therefore were included in downstream analyses (Supplemental Figure S2A: cyan). These included 502 animals’ lifespans that were censored and 132 deaths. Versus the *F.anselli* data set, these data for *F.damarensis* were richer for young and sparser for old ages (Supplemental Figure S2A). Nonetheless, these two species exhibited similar demographic properties, both in terms of Kaplan-Meier (Supplemental Figure S2B) and age-specific mortality hazard (Supplemental Figure S2C). For additional downstream analyses, we combined the data from *F.anselli* and *F.damarensis* into a single data set, representing the *Fukomys* genus (Supplemental Figure S2A: green). The combined *Fukomys* data set included lifespan data from 1041 animals, 966 of which survived past T_sex_ (270 days) so were included in downstream analyses; these included 319 deaths and 647 censorships.

Kaplan-Meier survival analysis revealed a *Fukomys* median lifespan of 3,622 days (Figure 5A). Unlike the survival curve for *H.glaber*, the *Fukomys* survival curve approached zero before being depleted of animals, allowing for calculation of a maximal lifespan, as previously defined (95th percentile; Ruby et al, 2018) of 7,369 days (Figure 5A), or 20.2 years. Age-specific mortality hazard was calculated for *Fukomys*, and again unlike *H.glaber*, substantial increase was observed with age (Figure 5B). Though less statistically robust, this trend was consistent with estimates taken from either the *F.damarensis* data alone, or from the *F.anselli* data alone, with or without accidental/fighting death censored (Supplemental Figure S2C).

**Fig 5:**
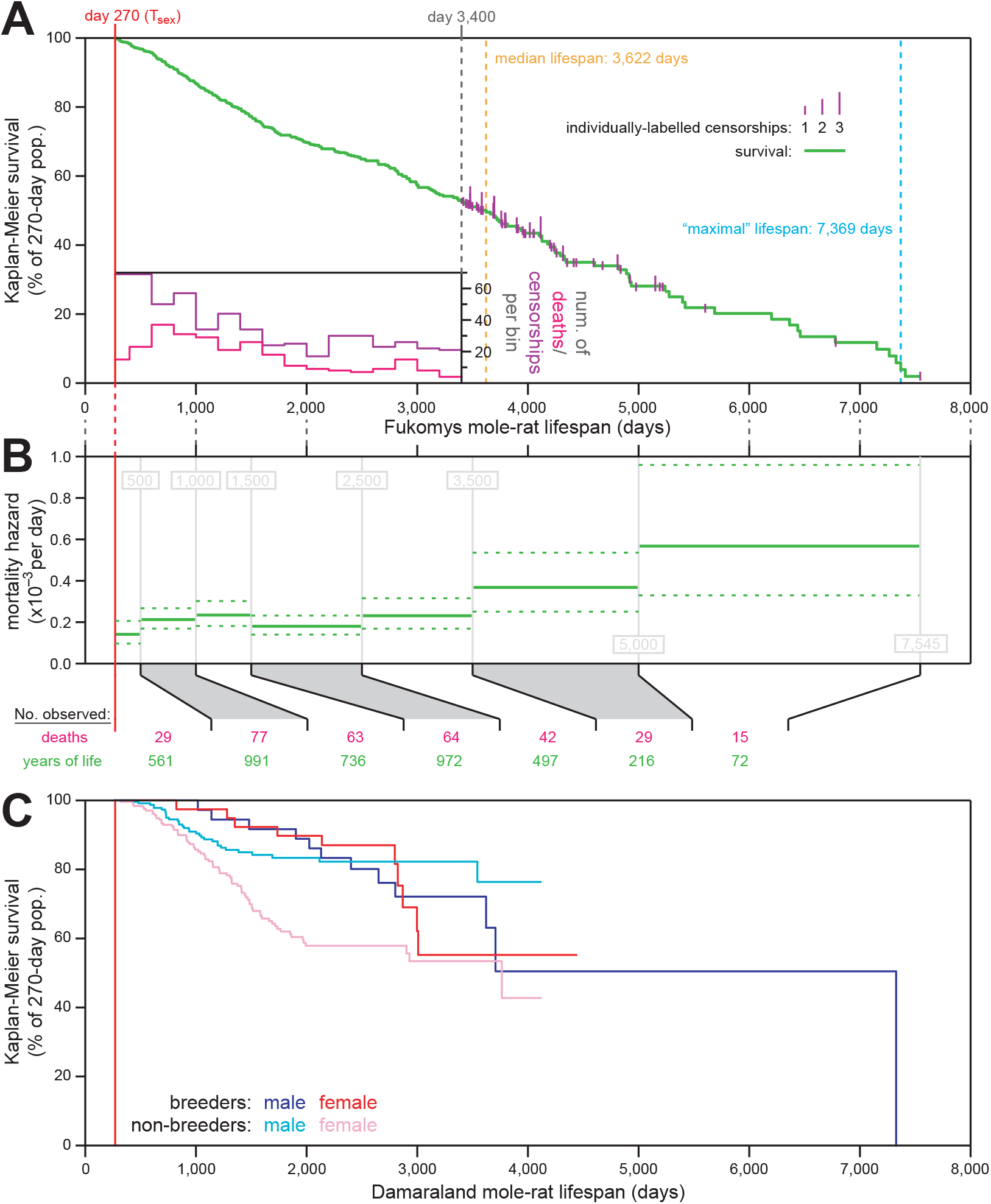
The mortality hazard of *Fukomys* mole-rats increased with age. (A) Kaplan-Meier survival curve for combined data from the *F.anselli and F.damarensis* species of mole-rats (green) after reaching reproductive maturity (T_sex_; 9 months from birth; 270 days; red). Plotted as in Fig. 1A, with a histogram of death and censorship (pink and purple, respectively) in the inset, up until 3,400 days of age, at which point 120 animals remained in the population; and with censorship events after that indicated by vertical ticks (purple), the size of which is proportional to the number of animals censored at each day-of-life. Equivalent per-species analyses are provided in Supplemental Figure S2. (B) Mortality-hazard estimates (solid green) with 95% confidence intervals (dotted green) across the observed *Fukomys* mole-rat lifespan. Estimates were calculated across time intervals of increasing size (demarked by grey lines) to compensate for decreasing accuracy-per-unit-time as the population size decreased. The borders of the age bins are indicated in gray, and the number of deaths (pink) and total years of animal life observed (green) per bin are indicated at the bottom of the panel. Equivalent per-species analyses are provided in Supplemental Figure S2. (D) For *F.damarensis* only: Kaplan-Meier survival starting at T_sex_ (270 days; red) for each of four reproductive categories of Damaraland mole-rats: male breeders (navy), female breeders, (red), male non-breeders (cyan), and female non-breeders (pink).

### Mole-rat mortality hazards showed lesser increase with age versus other mammals, with *H.glaber* still uniquely constant

Age of reproductive maturity (T_sex_) for a species is also generally predictive of species MLSP (Hamilton, 1966; Rose et al., 2007; Prothero, 1993). As described in Ruby et al (2018) and shown again here (Figure 6A), the failure of naked mole-rat mortality hazard to increase with age, even at many-fold the species’ T_sex_, is unique among the mammals analyzed. These included laboratory mice (*Mus musculus*) using the raw data for the control mice cohort from the published paper of Miller (Miller et al., 2014), with T_sex_ at 42 days post-natal (Tacutu et al., 2018) and a median lifespan of 847 days; humans (*Homo sapiens*), as reported for a 1900 birth cohort (Bell and Miller, 2005) and with T_sex_ defined as 16 years (Apter, 1980; Anderson et al., 2003); and horses (*Equus ferus caballus*), using insurance tables (Valgren, 1945) and T_sex_ at three years (Tacutu et al., 2018). All of these species are anciently diverged from *H.glaber*, with its closest relative being *M.musculus*, that diverged from *H.glaber* ∼73 MYA (Kim et al., 2011). In contrast, *H.glaber* diverged from humans ∼88 MYA and from horses ∼95 MYA (Springer et al, 2003). These evolutionary relationships are depicted in Figure 6B.

**Fig. 6:**
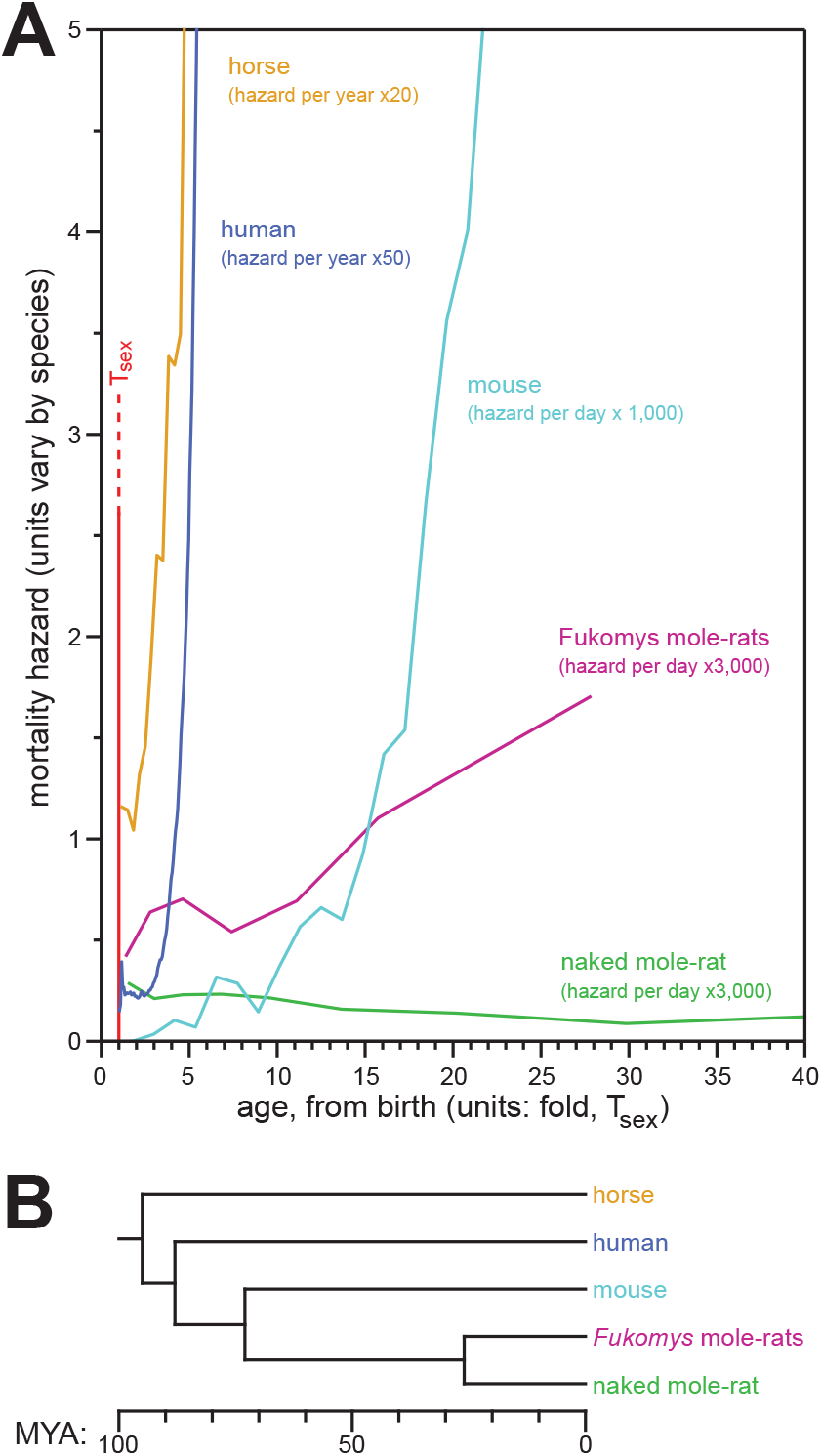
In contrast to the mortality hazards of other mammals, which increased with chronological age, the mortality hazard of naked mole-rats remained constant. (A) Age-specific mortality hazard for humans (navy), mice (cyan), horses (orange), *Fukomys* mole-rats (purple), and naked mole-rats (green); with age (x-axis) normalized to T_sex_ for each species. Reproduced from (Ruby et al, 2018), with *Fukomys* data added and *H.glaber* data updated using results from this study. See Results for references to original data sources. (B) A phylogenic tree for the species from panel (A). See Results for references to data sources.

Here, we added *Fukomys* mole-rat mortality hazard, age normalized to a T_sex_ of 270 days, to our comparative mammalian hazard plot (Figure 6A). *Fukomys* diverged from *H.glaber* 26 MYA (Figure 6B; Fang et al, 2014) and displayed an intermediate rate of hazard increase between the non-increasing hazard of *H.glaber* and the exponentially-increasing hazard of other mammals. *Fukomys* mortality hazard appeared to increase linearly, rather than with a Gompertzian exponential. The sparsity of late-life data for that genus made conclusions tentative. Additionally the uncertainty of age at T_sex_ made these conclusions tentative: we based our value of 270 days on our own experience, but alternative T_sex_ values are reported for *F.anselli* and *F.damarensis* on the AnAge website (Tacutu et al, 2018). However, with only 2.0% of the population surviving until the collection of the final datum (Figure 5A), it was unlikely that trend derived from observation through an insufficient portion of the genus’ lifespan.

## Discussion

### The elusion of demographic aging by *H.glaber* was replicated in an additional 5 years of new data

In this study, we re-visited our analyses and conclusions from (Ruby et al, 2018), bringing five years’ worth of additional data collection to bear. That period represented a period of unprecedented scale and stability for our collection: while animals continued to be killed for research (KFR) purposes, there were fewer constraints on husbandry resources to require culling of the population. Animals remained at the same institutional vivarium across that period (Calico Life Sciences, LLC), thus avoiding the potentially confounding husbandry challenges of institutional and geographic relocation. Especially given the mature status of the collection across this period (i.e., many animals of advanced age already present), these conditions facilitated an approximate doubling of the available observation numbers for demographic analysis across most age categories (Figure 2A).

These additional data provided two opportunities: first, to repeat our original analyses on the full, updated historical data, which added resolution and statistical confidence to our original claims (Figure 1). Second: the scale of data collected since our last study facilitated an independent analysis of *H.glaber* lifespan demography, performed entirely on data collected after the completion of data collection for our prior study (navy in Figure 2). Importantly, that independent analysis validated our original claims about naked mole-rat demography.

### Elevated hazard for non-breeders in small colonies

We previously reported that breeders of both sexes of naked mole-rats have substantially lower mortality hazards than non-breeders at all ages (Ruby et al, 2018), and that result was reproduced here (Figure 3). This result was not limited to *H.glaber*: our analysis of *F.damarensis* survival also revealed an advantage for breeders over non-breeders. This trait is shared with many cooperative breeding mammals (Sharp and Clutton-Brock, 2010; Dammann et al., 2011, Cram et al., 2018). These observations conflict with the disposable soma theory of aging (Kirkwood, 1977) but do not negate it: there are also multiple biological and sociological explanations for the survival advantage of socially-dominant individuals (e.g. Clarke and Faulkes, 1997; Medger et al., 2019; Sahm et al., 2021; Coen et al., 2021; Berger et al., 2018) that may confound one another and variably contribute to the fates of breeders versus non-breeders.

In order to gain deeper insight into the health consequences of naked mole-rat social hierarchy amongst non-breeders, we analyzed historical records of body weight and colony membership in terms of their impacts on mortality hazard (Figure 4). Dominance in naked mole-rat colonies is largely determined by aggressive pushing, tail-tugging, and shoving behaviors, so smaller animals are generally lower in the social hierarchy than larger (Clarke and Faulkes 1997; Toor et al., 2020; Holmes and Goldman, 2021). It was therefore unsurprising to observe that smaller adult animals had higher mortality hazard than bigger animals (Figure 4C), though that trend did not attain statistical significance in our analysis.

More surprising was the observation that non-breeders who were members of large colonies had lower mortality hazard than non-breeders who were members of small colonies, an observation that was statistically significant for both sexes (Figure 4E). One might have expected fewer colony members to result in less aggressive competition for, and therefore better access to, limited resources, e.g., food and heating pads (Clarke and Faulkes, 2001). However, such concerns may have been less relevant to our captive colonies than in the wild: animal husbandry rules necessitate the scaling up of all resources in parallel with colony growth (Smith & Buffenstein, 2021).

Further, even in the wild, intra-group competition is often trumped by cooperative advantages to produce overall survival advantages for members of larger social groups across a variety of species (Jarvis et al., 1998; Silk et al., 2010; Broom et al, 2009; Clutton-Brock, 2021). Larger colonies can also provide greater stability, with a well-established successfully breeding female and a cohort of older siblings having already resolved their dominance status and keeping the younger animals in check (Smith & Buffenstein, 2021).

### The mortality hazard of *Fukomys* mole-rats increased gradually with age

We sought to provide new evolutionary context for the exceptional longevity of naked mole-rats through demographic analysis of mole-rats from the *Fukomys* genus. Though substantially diverged from *H.glaber* 26 MYA (Fang et al., 2014), *Fukomys* mole-rats offered a phylogenitically closer comparator versus other mammals for which data are available (Figure 6B). Further, *Fukomys* mole-rats share the property of eusociality with naked mole-rats, which is thought to be connected to exceptional lifespan by evolutionary pressures (Kim et al., 2011; Fang et al., 2014). Demographic data were published for Ansell’s mole-rat, *F.anselli* (Dammann et al., 2019); we supplemented those data with lifespan data from the closely-related Damaraland mole-rat, *F.damarensis*. Those two species are estimated to have diverged ∼1-1.5 MYA (Van Daele et al, 2007), and they shared a similar ecological niche in the wild and statistically consistent life history patterns in captivity (Supp Figure S2). We combined the data from these species for a *Fukomys* genus-wide demography analysis and observed hazard to increase with age (Figure 5B).

While the increase of mortality hazard with age among *Fukomys* mole-rats contrasted with the age-independent hazard of *H.glaber*, it was gradual versus the Gompertzian (i.e. exponential) increase typically exhibited by mammals (Figure 6A). Various evolutionary theories highlight multiple aspects of mole-rat biology and lifestyle that could either drive or derive from the evolution of such extended longevity.

Subterranean rodents, regardless of clade or whether they are solitary or social, live longer than expected for their size, with environmental protection from extrinsic mortality (Buffenstein 2005; Healy et al., 2014). Similar protection from extrinsic mortality is also provided by herd protection/social lifestyle (Buffenstein 2005;Healy et al., 2015). Low extrinsic hazards give evolutionary tinkering with aging rates the opportunity to manifest selective advantage (Jacob, 1977), a premise backed up by several cross-species studies (Gaillard & Lemaître 2017). In the context of eusociality, kin selection and the maintenance of stable social hierarchies can provide further selective advantages to the extension of lifespan for breeders (Clarke and Faulkes, 1997; Medger et al., 2019; Sahm et al., 2021; Coen et al 2021). The converse may also be true, with enhanced longevity facilitating a eusocial lifestyle (Ross et al., 2015): increased longevity may facilitate the cohabiting of parents and multiple litters of offspring to form a eusocial colony, enhancing mutual beneficial opportunities for cooperative living and overall fitness of the social group (Carey, 2001; Lucas and Keller, 2020).

Regardless of which evolutionary theory best applies to the natural history of *H.glaber* and *Fukomys* mole-rats, their shared (though unequal) defiance of Gompertzian aging demonstrates that exponentially-increasing hazard with advancing age is not a fixed property of mammalian biology. While it seems to be the default, evolutionary mechanisms have managed to dampen or eliminate it when provided with the right selective incentives. It is not clear the extent to which both *H.glaber* and *Fukomys* benefit from longevity genes that arose in their common ancestor, versus evolved independent longevity-promoting mechanisms in the 26 million years since their divergence. However, their unequal successes suggest that multiple mechanisms are needed to counteract Gompertzian aging, and that not all mechanisms are at play in both lineages. The perspective that some age-slowing mechanisms are likely to be conserved between *H.glaber* and *Fukomys*, and that others are likely to be specific to *H.glaber*, should help the comparative biology community make maximal use of these two valuable natural resources for the science of aging.

## Materials and Methods

### Animal husbandry

Husbandry practices for *H.glaber* were identical to those described in (Ruby et al, 2018). A similar husbandry protocol was used for the Damaraland mole-rats.

The original progenitors of our Damaraland mole-rat colony were trapped near the town of Hotazel, South Africa (27°58′S, 17°41′E), and near the town of Dordabis in Namibia (22°58′S, 17°41′E). Like the naked mole-rat, all animals were microchipped at 90 days with a unique RFID number. Similarly, Damaraland mole-rats were group-housed in interconnected transparent multi-cage systems, joined together by plexiglass clear tubing in a climatically controlled room maintained at 26-28°C, 21% oxygen and with RH ranging between 20-50% and with a 12L:12D light cycle. Animals were fed daily a variety of fresh fruit and vegetables, including apples, yams, carrots, pumpkin, and other seasonal vegetables, replacing the natural geophytes upon which they feed. No water was provided as the animals meet all their water requirements through their diet or metabolic water production.

### *H.glaber* demographic data

Demographic analysis of *H.glaber* was performed on a newly-compiled aggregation of lifespan data, from across the collection of naked mole-rats described in (Ruby et al, 2018), which had been maintained and expanded under the same husbandry conditions described there. For this study, a snapshot of records was compiled on March 15, 2021, at which point they contained information for 7,536 animals from across the full history of the collection. Of those, complete birth and death/right-censorship information (single-day resolution), were available for 6,949 animals. For those animals, and versus our prior snapshot of records from this collection – provided as “Supplementary file 1” in (Ruby et al, 2018) – 3,329 could be matched to the prior records. Of those, 2,806 had perfect matches to RFID tags, birth dates, sex, breeder status, colony name, and logically consistent death/censorship data (for animals alive at the prior snapshot, new death/censorship dates from after those dates; otherwise, perfect matches). For an additional 94 animals, all data listed above were identical except for either colony name or breeding status: both of these parameters are expected to change for animals moved out of a colony to establish a new colony as a founding breeder. Combined, 3,222 animals matched for all vital data (RFID, birth date, sex, and consistent death/censorship). For an additional 94 animals, one of the four vital data had been updated: 11 for RFID (consistent with occasional replacement of RFID chips, or placement of RFID chips in previously un-chipped animals); 32 for sex (consistent with inaccurate initial sex determination for young animals); 40 for date of birth (consistent with review of birth data at the time of RFID implantation); and 11 with updated and inconsistent death/censorship dates. For all 94 of those animals, the updated values were used for analysis. For an additional five animals, two pieces of vital data differed: one had inconsistent death/censorship and sex; one had inconsistent death/censorship and date of birth; and three had different dates of birth and sexes. For those animals as well, the updated values were used for analysis. For an additional eight animals, exact dates of death/censorship were missing from current records but found in the prior snapshots, with all other vital data matching. For those animals, the death/censorship values and dates from the (Ruby et al, 2018) snapshot were used. An additional 26 animals had RFID matches to animals that had missing or low-resolution birth and/or death/censorship data in the (Ruby et al, 2018) snapshot; those animals were assigned their original table ID’s and current data values. In total, 3,355 animals were paired to individuals cataloged in “Supplementary file 1” of (Ruby et al, 2018): all of those are presented in Supplemental Data File 1 using their original ID’s (e.g. “Hg-02241”).

An additional 3,602 animals with day-resolution birth and death/censorship data were newly added to the roster presented in Supplemental Data File 1, versus the catalog from (Ruby et al, 2018). Those animals were all assigned ID numbers ≥5000 (e.g. “Hg-05072”). The vast majority of those animals were born after (3,244) or during (33) the prior data consolidation period (April 14 – May 17, 2016). Of the remaining 325 animals, the large majority (285) were born within a year prior to April 14, 2016, reflecting a normal lag for husbandry records to be entered into our database.

In summary: 6,949 naked mole-rats were cataloged with day-resolution birth and death/censorship data, 3,355 of which could be matched to animals from (Ruby et al, 2018). An additional eight animals with missing death/censorship data in the current inventory were included using those data from (Ruby et al, 2018), bringing the total number of animals analyzed to 6,957. For still-surviving animals, right censorship was applied at the date of record aggregation: March 15, 2021.

### Analysis of lifespan and age-specific mortality hazard

Except as otherwise specified in the text, right censorship status was applied to animal lifespans in three cases: if the animal was (1) transferred to another collection, with the date of transfer considered as the censorship event; (2) euthanized for research purposes when still healthy, with the date of sacrifice as the censorship event; or (3) still alive when the process of compiling records was completed, with the date of completion as the censorship event. Left censorship was applied to entire data sets as specified in the text, generally at specific historical dates in order to exclude data collected prior to that date.

Kaplan-Meier survival was calculated using their method (Kaplan and Meier, 1958). Mortality hazards were calculated for each specified age bin/group of animals as the number of observed death events (i.e., number of days on which a naked mole-rat was observed to have died) divided by the total number of days that naked mole-rats were under observation, limited to that age bin/population. Confidence intervals were calculated using the Wilson score interval with continuity correction (Newcombe, 1998).

### *H.glaber* body weight analysis

Historical records of body weights, collected in the course of animal health checks, were aggregated concurrently with the lifespan data from (Ruby et al, 2018). These records were less complete than the demographic data, but nonetheless included 19,118 measurements taken from 1,940 animals with complete lifespan data. For most analyses, only weights taken after 500 years of age were used: there were 6,785 such measurements, taken from 942 animals. The demographic data used for these analyses was as compiled for (Ruby et al, 2018), and it is provided along with colony membership and body weight data in Supplemental Data File 2. Animals from (Ruby et al, 2018) with full demographic data but no body weight data are also included in that file. Animal ID’s in that file are matched to (Ruby et al, 2018) and Supplemental Data File 1.

For mortality versus body weight, analysis was performed for survival across a single year. For each animal with body weight(s) measured after 500 days of age, survival data were analyzed starting from the date of the last recorded body weight (i.e. left-censored on that date). If the animal survived uncensored for over a year, then right-censorship was applied one year after the weight measurement. Animals with qualifying body weight values were sorted into quartiles based on those final measurements, and mortality hazard was estimated across all of the animals in each quartile. Confidence intervals were calculated using the Wilson score interval with continuity correction (Newcombe, 1998).

### *H.glaber* colony size analysis

For mortality versus colony size, analyses were performed of survival across a single year. The results for each year are shown in Supplemental Figure S1: for years 2012-2015, demographic records from Ruby et al (2018; also provided in Supplemental Data File 2) were used, and April 14 was used as the cut-off date; for years 2016-2020, the updated demographic records from this manuscript were used (Supplemental Data File 1), and March 15 was used as the cut-off date. Cut-off dates were selected to match the data compilation date for each data set, ensuring a full year’s worth of data availability for each year’s analysis. The shift to an older record compilation for earlier-year analyses was to minimize the effect of movement out of colonies as a confounder: colony affiliations for each compiled data set reflected colony membership at the time of compilation and did not include prior membership history. Importantly, this caveat should have had minimal impact: due to violence against intruders, animals are never introduced into existing colonies, only removed from colonies in order to establish new colonies as breeders (Smith and Buffenstein, 2021), and breeders were excluded from these analyses.

For analysis of colony sizes, only animals at least 500 days old by each cut-off date were considered. This decision was justified by animals younger than that age being not fully grown and therefore potentially presenting less competitive rivalry; and by the inconsistent integration of animals much younger than one year (pre-chipped) old into our demography database. But it may have added variance in the form of colony-size error in the cases of colonies with highly skewed age distributions. Analyses were also conducted in a sex-specific manner, i.e., considering only the counts of same-sex, non-breeding animals for colony size. This decision was justified under the assumption that competition and hierarchy may be different between the two sexes (Reeve & Sherman, 1991), but may have added variance in cases where the sex ratio was skewed in a colony, to the extent that social hierarchy among non-breeders is sex-independent.

For each cut-off date, for each sex, all qualifying animals were used to generate a census (animal count) for each represented colony name. Individual animals were sorted into quartiles based on the sizes of the colonies of which they were members; tied animals were assigned to adjacent quartiles arbitrarily. For members of each quartile, the mean and standard deviation of colony sizes for each animal were calculated, and the mortality hazard for those animals was calculated across one year, beginning at the cut-off date (i.e. left-censored on the cut-off date, and right censored one year after the cut-off date for animals that were not deceased or otherwise censored during the year). Confidence intervals were calculated using the Wilson score interval with continuity correction (Newcombe, 1998).

For regression of mean colony size versus mortality hazard, for each sex: for all quartiles across all years shown in Supplemental Figure S1, mean colony sizes were treated as x-axis values and mortality hazard estimates as y-axis values, and linear regression was performed using the “scipy.stats.linregress” function from Scipy (Virtanen et al., 2020), which in addition to slope and intercept returns Pearson R and regression p-value.

### *F.damarensis* demographic data

As for *H.glaber*, lifespan data from our longstanding collection of Damaraland mole-rats, sp. *F.damarensis*, were compiled on March 15, 2021, with that date used as the right-censorship point for still-living animals. Of 754 animals with records, 702 had day-resolution birth and death/censorship data and were therefore included in these analyses. Per-animal demographic data are provided in Supplemental Data File 3.

### Calculation of maximum versus observed expected lifespan

Allometric expectations were determined using the equation from (de Magalhães et al., 2007):

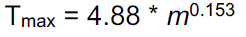

where body mass *m* is in grams and T_max,_ predicted maximum lifespan, is in years. The longevity quotient (LQ) was defined as the ratio of observed MLSP to T_max_. For *H.glaber*: using a body mass = 35g, T_max_ = 8.4 years; and with observed MLSP = 37 years, LQ = 4.4. For *F.anselli*: using a body mass = 90g, T_max_ = 9.7 years; and with observed MLSP = 22 years, LQ = 2.3. For *F.damarensis*: using a body mass = 160g, T_max_ = 10.6 years; and with observed MLSP = 20 years, LQ = 1.9.

## Supporting information

Supplemental raw data

## Acknowledgements

We thank the numerous animal attendants who have diligently cared for the naked mole-rats and Damaraland mole-rats over the past three-and-a-half decades.

## Ethics

Animal experimentation: All of the animals were handled according to approved institutional animal care and use committee (IACUC) protocols, most recently of the Buck Institute (A10138). IACUC approval for housing these colonies of naked mole-rats and Damaraland mole-rats were also obtained from all institutions at which these collections were historically housed.

## Supplemental Files

**Supplemental Figures:** two Supplemental Figures (S1 & S2), with legends, as one PDF.

**Supplemental Data File 1**:

Full *H.glaber* lifespan data, compiled in 2021 (tab-separated text file).

**Supplemental Data File 2**:

Full *H.glaber* lifespan data, as compiled in 2016, with body weight measurements (tab-separated text file).

**Supplemental Data File 3**:

Full *F.damarensis* lifespan data, compiled in 2021 (tab-separated text file).

**Supplemental Data File 4**:

Wherever practical: numerical values that are plotted across all Figures and Supplemental Figures, organized by Figure (Excel notebook file).

