## Supplementary figures and images for "Five years later, with double the demographic data, naked mole-rat mortality rates continue to defy Gompertzian laws by not increasing with age"

### Supplementary Figure 1

Supplemental Figure S1

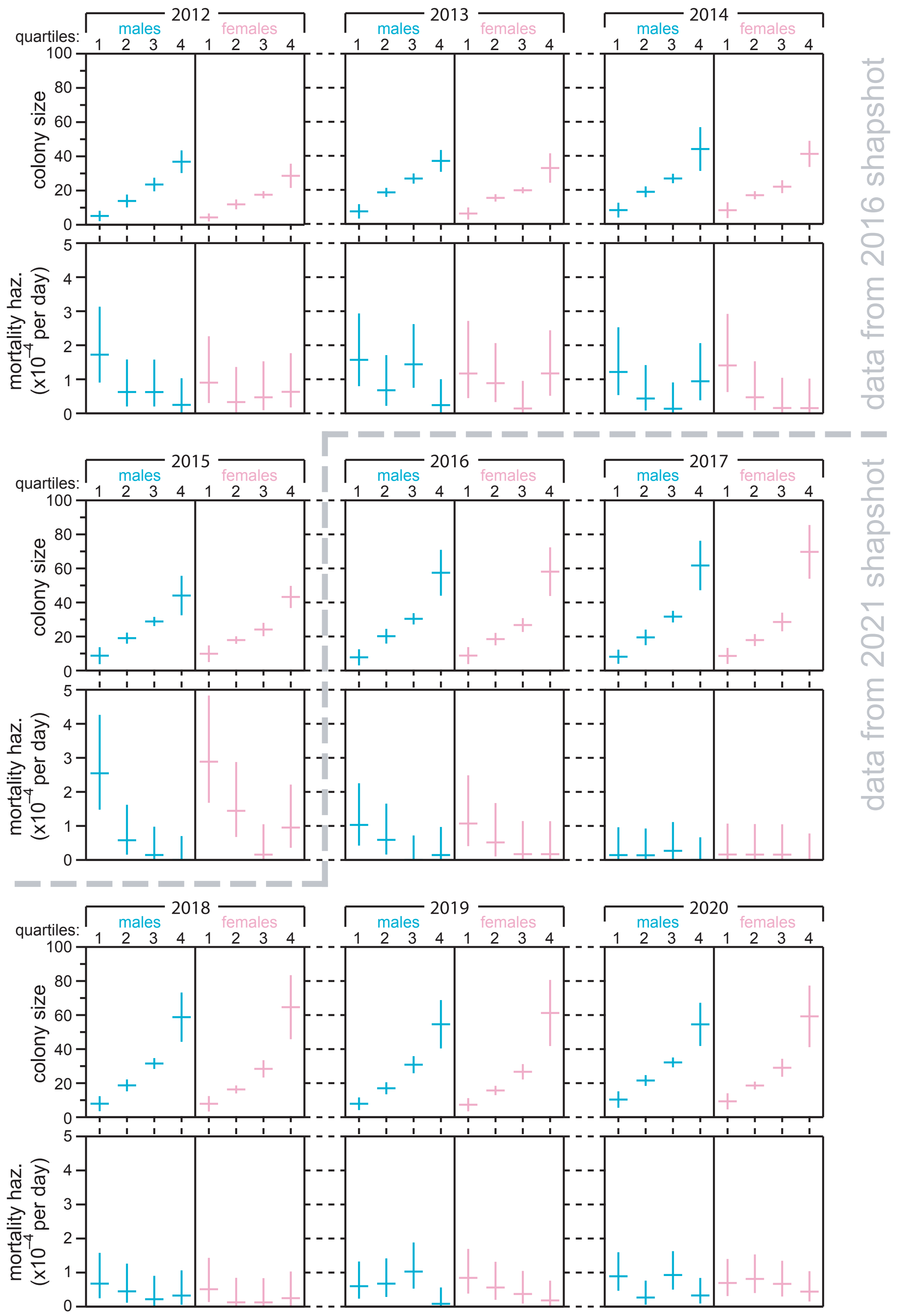

### Supplementary Figure 2

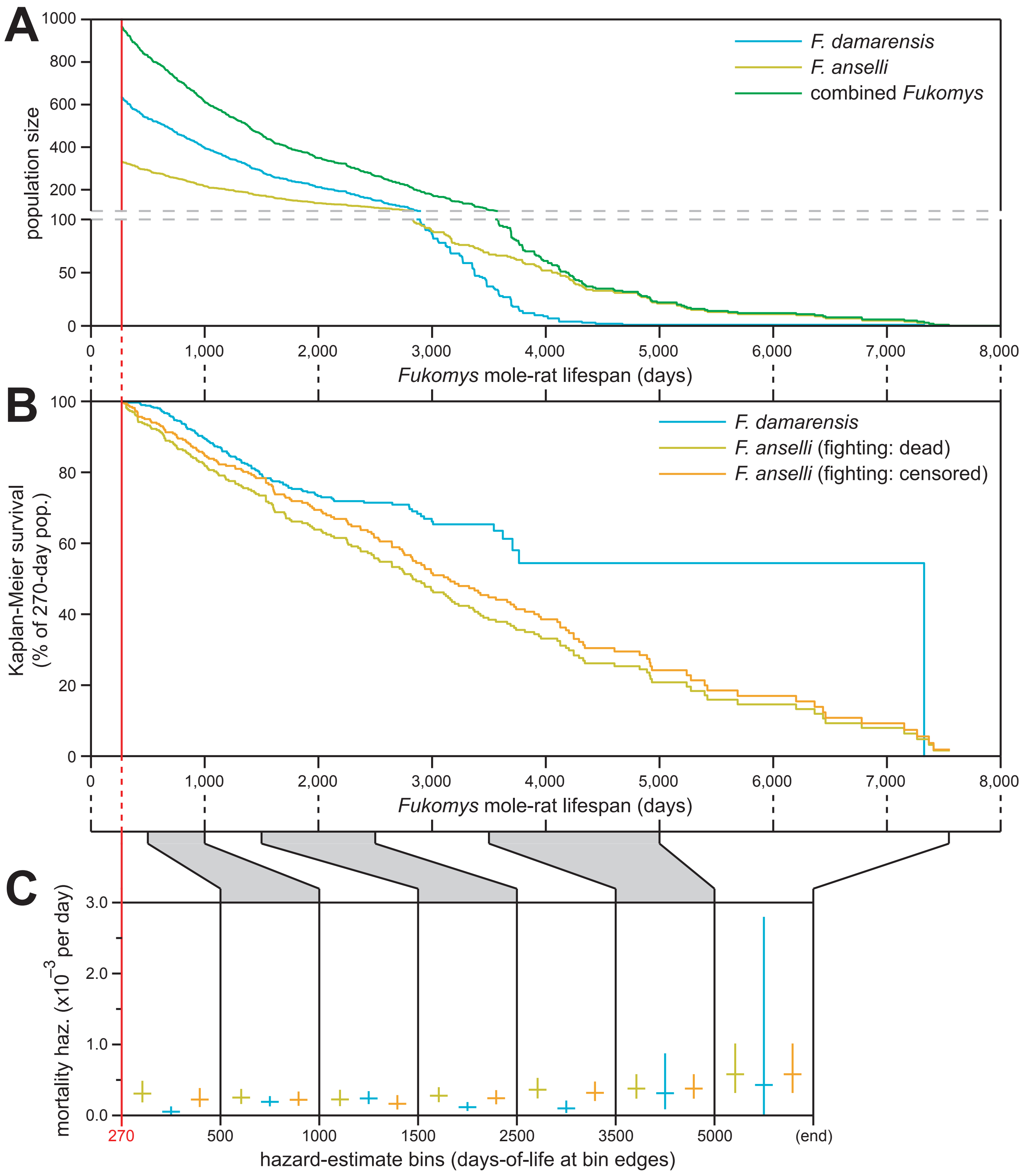
